# Stability of SARS-CoV-2 in cold-chain transportation environments and the efficacy of disinfection measures

**DOI:** 10.1101/2022.08.09.503429

**Authors:** Shuyi Peng, Guojie Li, Janak L. Pathak, Xiaolan Guo, Yuyin Lin, Hao Xu, Wenxi Qiu, Jiaying Zheng, Wei Sun, Xiaodong Hu, Guohua Zhang, Bing Li, Xinhui Bi, Jianwei Dai

## Abstract

Cold-chain environment could extend the survival duration of SARS-CoV-2 and increases the risk of transmission. However, the effect of clod-chain environmental factors and packaging materials on SARS-CoV-2 stability and the efficacy of intervention measures to inactivate SARS-CoV-2 under cold-chain environment remains uncertain. This study aimed to unravel cold-chain environmental factors that preserved the stability of SARS-CoV-2 and disinfection measures against SARS-CoV-2 under the cold-chain environment. The spike gene of SARS-CoV-2 isolated from Wuhan hu-1 was used to construct the SARS-CoV-2 pseudovirus and used as model of the SARS-CoV-2 virus. The decay rate of SARS-CoV-2 pseudovirus in the cold-chain environment, various types of packaging material surfaces i.e., PE plastic, stainless steel, Teflon and cardboard, and in frozen seawater was investigated. The influence of LED visible light(wavelength 450 nm-780 nm) and airflow movement on the stability of SARS-CoV-2 pseudovirus at -18° C were subsequently assessed. The results show that SARS-CoV-2 pseudovirus decayed more rapidly on porous cardboard surface compared with the non-porous surfaces including PE plastic, stainless steel and Teflon. Compared with 25° C, the decay rate of SARS-CoV-2 pseudovirus was significantly lower at low temperature. Seawater preserved viral stability both at -18° C and repeated freeze-thawing cycles compared with deionized water. LED visible light illumination and airflow movement environment at -18° C reduced the SARS-CoV-2 pseudovirus stability. In conclusion, our results indicate cold-chain temperature and seawater as risk factors for SARS-CoV-2 transmission and LED visible light illumination and airflow movement as possible disinfection measures of SARS-CoV-2 under the cold-chain environment.

**Importance:** It is widely recognized that low temperature is a condition for maintaining virus vitality, and cold-chain transportation spreads the events of the SARS-CoV-2 were reported. This study provides that the decay rate of the SARS-CoV-2 pseudovirus at low temperatures varies on different packaging materials, and salt ions present in frozen foods such as seafood may protect virus survival. These results provide evidence for the possibility of SARS-CoV-2 transmission through cold-chain transport and also suggest the importance for disinfection of items. However, the commonly used disinfection methods of ultraviolet radiation and chemical reagents are generally not suitable for the disinfection of frozen food. Our study shows LED visible light illumination and airflow movement as possible disinfection measures of SARS-CoV-2 under the cold-chain environment. This has implications for reducing the long-distance transmission of the virus through cold-chain transportation.

## 1. Introduction

Severe acute respiratory syndrome coronavirus 2 (SARS-CoV-2) is the causative agent of coronavirus disease 2019 (COVID-19) and the seventh coronavirus documented to infect humans. An unparalleled pandemic of COVID-19 has created a worldwide threat to public health. As of May 2022, the total number of confirmed cases of COVID-19 was over 50 million, with more than 6 million deaths globally (WHO,2022). The COVID-19 resurgence due to contaminated imported food or packaging via cold-chain logistics, where the SARS-CoV-2 was found not only on the outer packaging but also in the inner packaging of the frozen food. On August 12, 2020, local authorities in Shenzhen, Guangdong Province of China had detected SARS-CoV-2 nucleic acid positive on the contaminated surface of the frozen chicken wing originated from the Brazil, which was the first found SARS-CoV-2 nucleic acid on the surface of the frozen food in China (Han et al., 2020; Huilai Ma1 and Ha, 2020; Pang et al., 2020). During the recurrence of SARS-CoV-2 in Qingdao, the infectious SARS-CoV-2 has been isolated from the outer surface of frozen cod, which was the first known case that the infectious SARS-CoV-2 was discovered in the cold-chain environment in the world. These events strongly supported the perspective that the cold-chain environment could be the risk factor to maintain infection and transmission of SARS-CoV-2. (Chen et al., 2022; He et al., 2021; Huilai Ma1 and Ha, 2020; Liu et al., 2020). Therefore, the transmission of the SARS-CoV-2 via cold-chain should not be neglected. Firstly, environmental conditions, such as temperature, have been shown to influence the decay rate of infectious viruses (Aboubakr et al., 2021). SARS-CoV-2 is sensitive to heat but can survive for more than 14 days at 4 ° C. Fisher et al. found that the titer of SARS-CoV-2 remained virus infectivity for over 21days in both refrigerated and frozen conditions (Chin et al., 2020; Fisher et al., 2021; Kumar et al., 2021b). All these findings suggest that maintained infectiousness (days to perhaps weeks) of SARS-CoV-2 under a cold-chain environment could be associated with the virus stability. However, there is a lack of information on the decay rate of SARS-CoV-2 under different temperature conditions. Secondly, cold-chain goods had become the medium for SARS-CoV-2 transmission of the virus, which is unrealistic for prevention and transmission control by interrupting the international trade of cold-chain for a long time (He et al., 2021). Thus, the question of how to reduce the risk of SARS-CoV-2 transmission in the cold-chain industry has become a challenge. Common disinfections including sodium hypochlorite and peroxyacetic acid can be easily frozen after being sprayed under the cold-chain environment, resulting in a sharp decline in disinfection efficacy. Meantime, the ultraviolet and ionization radiation methods may affect the flavor and texture of goods and increase the risk for both humans and the environment (Li et al., 2020; Wu et al., 2022; Zucker et al., 2021). Therefore, it is vital for exploring effective methods to inactivate SARS-CoV-2 under cold-chain environment.

The key purpose of the present study was to explore the stability of SARS-CoV-2 under the cold-chain environment and to test the efficacy of physical methods for the disinfection of SARS-CoV-2. Thus, we constructed SARS-CoV-2 pseudovirus model containing the SARS-CoV-2 spike glycoprotein. Recent studies had been demonstrated that the protruding spike proteins of SARS-CoV-2 catalyze interactions with surfaces and mediate the virus entry into the host cells (Yao et al., 2020). Therefore, SARS-CoV-2 pseudovirus was employed as a model to investigate the stability of the SARS-CoV-2 under the cold-chain based on the reports from literature (Liu et al., 2021; Peinetti et al., 2021; Yi et al., 2020; Zucker et al., 2021). In addition, physical methods have been proposed for the inactivation of SARS-CoV-2 in the cold-chain environment to minimize the risk of the cold chain-related SARS-CoV-2 transmission.

## 2. Materials and methods

### 2.1 Construction and infection with SARS-CoV-2 pseudovirus

SARS-CoV-2 pseudovirus with spike glycoprotein was constructed as previously described (Neerukonda et al., 2021). The human embryonic kidney cells (HEK293T) and HEK293T cells with stable expression of human ACE2 (293T-hACE-2 cells) were provided by Weisheng Guo’s Lab. The cells were maintained in Dulbecco’s modified Eagle’s medium (DMEM, Gibco) with 10% FBS (Gibco), penicillin (100 U/mL) and streptomycin (100 mg/mL). All the cells were incubated in 5% CO2 incubator at 37° C. Briefly, HEK293T cells grown to 80% confluency were co-transfected with pcDNA3.1-SARS-CoV-2-S (Genewiz Biotech Co., Ltd., Suzhou, China), pLV-EGFP-luciferase-N and psPAX2 plasmid using polyethyleneimine (PEI, Polysciences, USA). Supernatants were harvested at 48 h and 72 h, concentrated using ultra-high-speed centrifugation (XPN-100, Beckman, USA), and stored at -80° C until use. The SARS-CoV-2 pseudovirus was added to 293T-hACE-2 cells. The viral titer was estimated by quantification of viral copy number in infected cells through the amplification of pseudovirus-specific transgene (Long terminal repeat, LTR) and a single copy reference gene (Albumin, ALB) using a real-time quantitative PCR assay (SYBR® Green Premix Pro Taq HS qPCR Kit (Accurate Biology Co. Ltd., Changsha, China) according to the protocol of the previous report (Barczak et al., 2015). The primers that were used for RT-qPCR are as follows. ALB-F: 5’
s-TTTGCAGATGTCAGTGAAAGAGA-3’, ALB-R: 5’-TGGGGAGGCTATAGAAAATAAGG-3’; LTR-F: 5’-CTAGCTCACTCCCAACGAAGA-3’, LTR-R: 5’-GGTCTGAGGGATCTCTAGTT-3’.

### 2.2 Transmission electron microscopy (TEM) assay

The SARS-CoV-2 pseudovirus was filtrated through a 0.45-micron pore-size membrane and concentrated by centrifuging for 2 h at 25000 g. The supernatant was removed and the virus was resuspended in ultrapure water containing 0.005% Bovine serum albumin (Sigma-Aldrich, USA). Purified and filtrated SARS-CoV-2 pseudovirus was firstly fixed in 2% glutaraldehyde (Leagene, Beijing, China), and then was negative-stained through 1% Phosphotungstic acid (Solarbio, Beijing, China). Finally, the SARS-CoV-2 pseudovirus samples were collected on a copper grid (Zhongjingkeyi, Beijing, China). The samples were observed using a transmission electron microscope (JEM-1400PLUS, Jeol Ltd., Akishima-shi, Japan).

### 2.3 Stability of SARS-CoV-2 pseudovirus under different surfaces

The indicated titer of SARS-CoV-2 pseudovirus (C_0_) was placed on the four types of common surfaces, including Teflon, cardboard, stainless steel, and PE plastic, for 3 days at 16° C and 70% relative humidity, which is similar to the outbreak of COVID-19 in Guangzhou. SARS-CoV-2 pseudoviruses were recovered using DMEM after being exposed to different surfaces and further infected to 293T-hACE2 cells to quantify the titer (C_t_) of pseudovirus as described above. Then, decay curves over time and decay rate constants *k* of SARS-CoV-2 pseudovirus were analyzed as previously reported (Ahmed et al., 2020).

### 2.4 Stability of SARS-CoV-2 pseudovirus under cold-chain temperature and in seawater at -18° C and repeated freeze-thawing cycles

The indicated titer of SARS-CoV-2 pseudovirus (C_0_) was placed under 25° C, 4° C, 0° C, -18° C, and -70° C for 20 days and suspended in seawater, which was collected from the sea of Hainan province, and in deionized water. The resuspended solutions were deposited at -18° C on the surface of the PE plastic respectively, and further maintained for 8 times freeze-thaw cycles treatment, with the temperature ranging from -18° C to 25° C. SARS-CoV-2 pseudoviruses were recovered and cultured with 293T-hACE2 cells to quantify the titer (C_t_) as described above. Subsequently, the decay curves over time and the decay rate constants *k* of the virus were analyzed as described above.

### 2.5 The disinfection potential of LED visible light illumination and airflow movement on SARS-CoV-2 pseudovirus under cold-chain conditions

The standard LED visible light containing spectral wavelengths of 450 nm to 780 nm, the luminous intensity of 4 mW/cm^2^, and the airflow movement with an airflow of 3m/s were used to treat SARS-CoV-2 pseudovirus at -18° C condition on the surface of packaging materials commonly used for marine products, including PE plastic and corrugated carton, which are smoother than cardboard. SARS-CoV-2 pseudoviruses were recovered and cultured with 293T-hACE2 cells to quantify the titer (C_t_) as described above. Subsequently, the decay curves over time and the decay rate constants *k* of the virus were analyzed as described above.

### 2.6 SARS-CoV-2 pseudovirus RNA damage assay

Total RNA of SARS-CoV-2 pseudovirus that was exposed under the LED visible light for 10 min was extracted using Steady Pure Virus DNA/RNA Extraction Kit (Accurate Biology Co. Ltd., Changsha, China), and further evaluated RNA damage using agarose gel electrophoresis. The gray-scale scanning analysis of pseudoviruses genome RNA degradation was performed using Image J software (Java-based image-processing and analysis software, National Institutes of Health, USA).

### 2.7 Decay rate calculations

In the linear regression analyses, the observed viable titer of SARS-CoV-2 pseudovirus concentrations was linearized using the natural log (ln)-transformation of the normalized concentrations as shown in equation (1). The first-order decay rate constant of pseudovirus at each point time was calculated by linear regression using GraphPad Prism Version 8.3.1 (GraphPad Software, La Jolla, CA, USA). As a comparison metric, *r*^2^ was used to evaluate the proportion of the variance explained by the linear model.

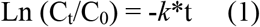

### 2.8 Statistical analysis

The data are presented as the mean ± standard deviation from three independent experiments. All statistical analyses were performed using GraphPad Prism 8.3.1 (GraphPad Software, La Jolla, CA, USA). One-way analysis of variance (ANOVA) with Bonferroni post-test was used for pairwise comparisons of multiple groups. The statistically significant difference between the two groups was considered while *p*<0.05.

## 3. Results

### 3.1 Construction and properties of SARS-CoV-2 pseudovirus

To investigate the infectivity and stability of SARS-CoV-2 under the cold-chain environment, SARS-CoV-2 pseudovirus particles were constructed using a lentivirus packaging system. Firstly, the SARS-CoV-2 pseudovirus was identified under TEM and showed that the lentivirus and pseudovirus were both 100∼120 nm in diameter spherical-shaped enveloped virus consisting of a lipid bilayer membrane and SARS-CoV-2 pseudovirus expressed spike glycoprotein protruding from the virion envelope membrane (Fig.1A). The infectivity of SARS-CoV-2 is related to the expression level of ACE2 (the receptor of SARS CoV-2) in the target cells.SARS-CoV-2 pseudovirus showed much higher infectivity in 293T-hACE-2 cells than in the regular HEK293T cells (Fig.1B). The results showed that the size, spherical-shaped enveloped virus with the spike-like crown structure wrapping around the pseudovirus membrane, and the infectiousness to 293T-hACE-2 cells of pseudovirus particles was consistent with the characteristics of SARS-CoV-2. Hence, SARS-CoV-2 pseudovirus was used as the ideal mimic to investigate the stability and infectivity of the SARS-CoV-2.

**Figure 1.**
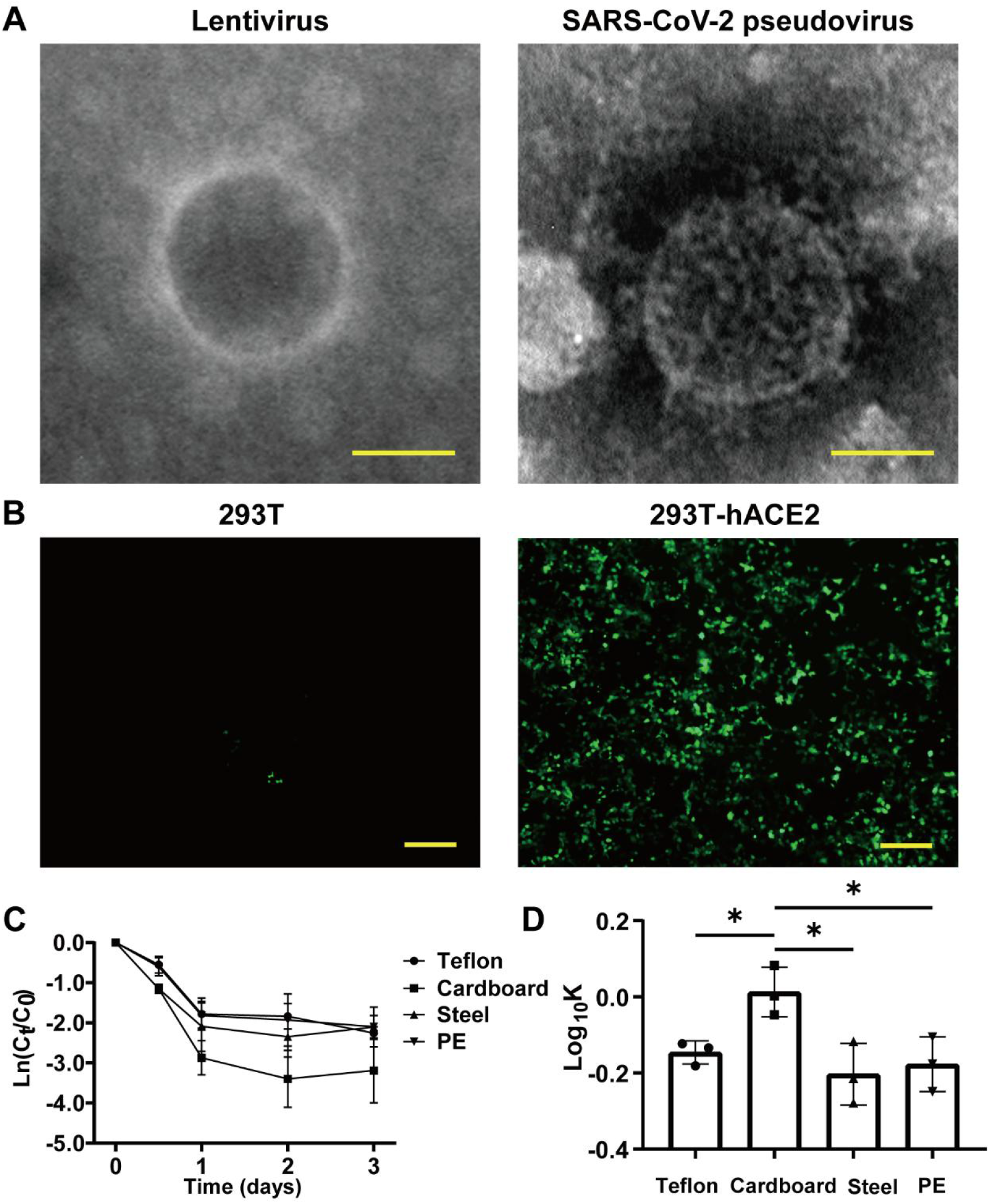
Identification of SARS-CoV-2 pseudovirus. (A) Structure analysis of lentivirus and SARS-CoV-2 pseudovirus using Transmission electron microscopy (TEM) assay. (B) Infectivity of SARS-CoV-2 pseudovirus on host cells. (C) Decay curves of SARS-CoV-2 pseudovirus over 3 days on the surface of Teflon film, cardboard, stainless steel, and PE plastic. (D) Log10-transformed mean pseudovirus decay rate constants *k* of SARS-CoV-2 pseudovirus on the surfaces of different materials at 16° C. Scale bar: 50 nm (A) and 200 μm (B). *, *p*<0.05.

To determine the mimic properties of the SARS-CoV-2 pseudovirus to SARS-CoV-2, we investigated the stability of SARS-CoV-2 pseudovirus on the different surfaces, including the cardboard, polyethylene (PE) plastic, stainless steel, and Teflon film. The decay of SARS-CoV-2 pseudovirus on the surfaces was analyzed using linear regression. The declining concentration and the mean decay rate constants *k* for SARS-CoV-2 pseudovirus on the different surfaces are shown in Fig.1C and Fig.1D. The SARS-CoV-2 pseudovirus was stable on the PE plastic, stainless steel, and Teflon and remained viable for three days on these surfaces. The mean decay rate revealed that the survival time of SARS-CoV-2 pseudovirus was no obvious differences on the non-porous surfaces; however, the survival time was surprisingly less on porous surfaces such as cardboard than on non-porous surfaces. For the infectivity of SARS-CoV-2 pseudovirus on the various surfaces, mean decay rate constants *k* ranged from 0.67/day at PE plastic to 1.04/day at the cardboard with *r*^2^ values from 0.65 to 0.67 (Table 1). The results indicate that SARS-CoV-2 pseudovirus remains stable for a longer period on the non-porous surfaces.

**Table 1.** The decay rate of SARS-CoV-2 pseudovirus on different container surfaces

| Surfaces | *K* (mean ± SD) | [95% CI slope] | R<sup>2</sup> |
| --- | --- | --- | --- |
| Teflon film | 0.715±0.118 | -0.957 to -0.474 | 0.76 |
| Cardboard | 1.037±0.213 | -1.50 to -0.576 | 0.65 |
| Stainless steel | 0.634±0.166 | -0.992 to -0.275 | 0.53 |
| PE Plastic | 0.670±0.132 | -0.955 to -0.386 | 0.67 |

NS: Non-significant deviation from the model; SD: standard deviation.

### 3.2 The decay of SARS-CoV-2 pseudovirus under the cold-chain environment

The sporadic outbreaks of COVID-19 had a high possibility to be related to the cold-chain environment. Hence, we detected the stability of SARS-CoV-2 pseudovirus under different temperature conditions at the PE surface, such as 25° C, 0° C, 4° C, -18° C, and -70° C for 20 days. The decay persistence of viable SARS-CoV-2 pseudovirus was in a temperature-dependent manner and the declining concentration of SARS-CoV-2 pseudovirus under room temperature was higher than that under other cold-chain temperatures and no SARS-CoV-2 pseudovirus was detected under room temperature at the sixth day (Fig. 2). In addition, SARS-CoV-2 pseudovirus could survive for over 20 days under cold-chain temperatures (Fig.2A). Accordingly, the decay rate constants *k* of SARS-CoV-2 pseudovirus under different cold-chain conditions were both significantly lower than that under room temperature (Fig.2B). For the stability of SARS-CoV-2 at various temperatures, mean decay rate constants *k* ranged from 0.08/day at -70° C to 0.87/day at room temperature with *r*^2^ values from 0.57 and 0.91 respectively (Table 2). These results indicated that cold-chain temperature conditions could decrease the decay rate of SARS-CoV-2 pseudovirus compared to the room temperature and increase the risk of SARS-CoV-2 transmission.

**Table 2.**
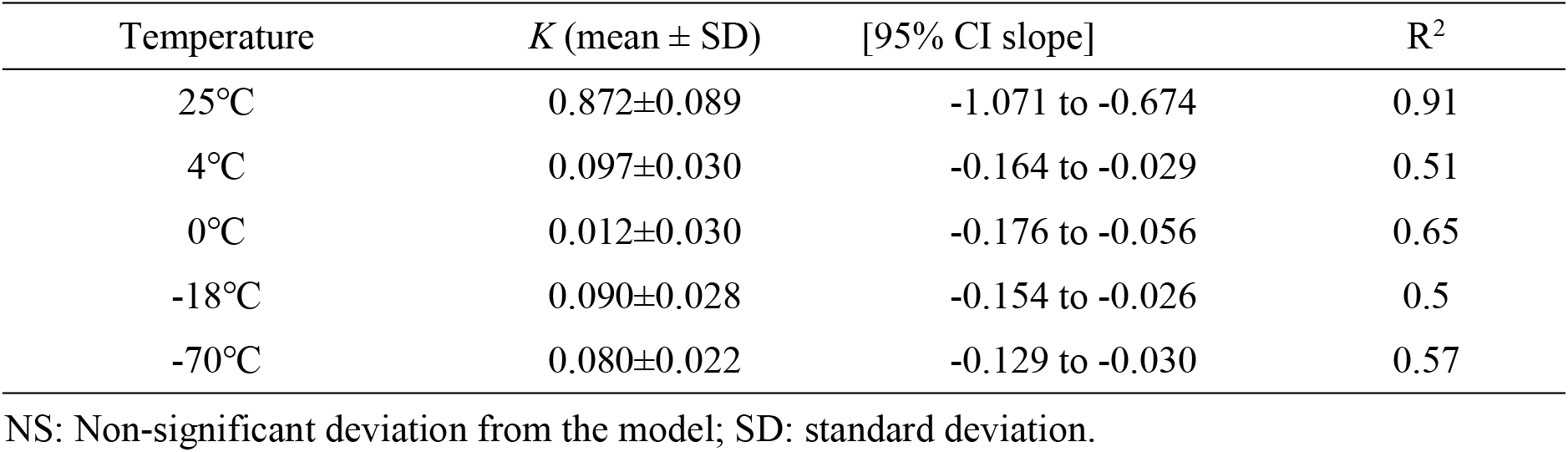
The decay rate of SARS-CoV-2 pseudovirus on cold-chain temperature conditions

**Figure 2.**
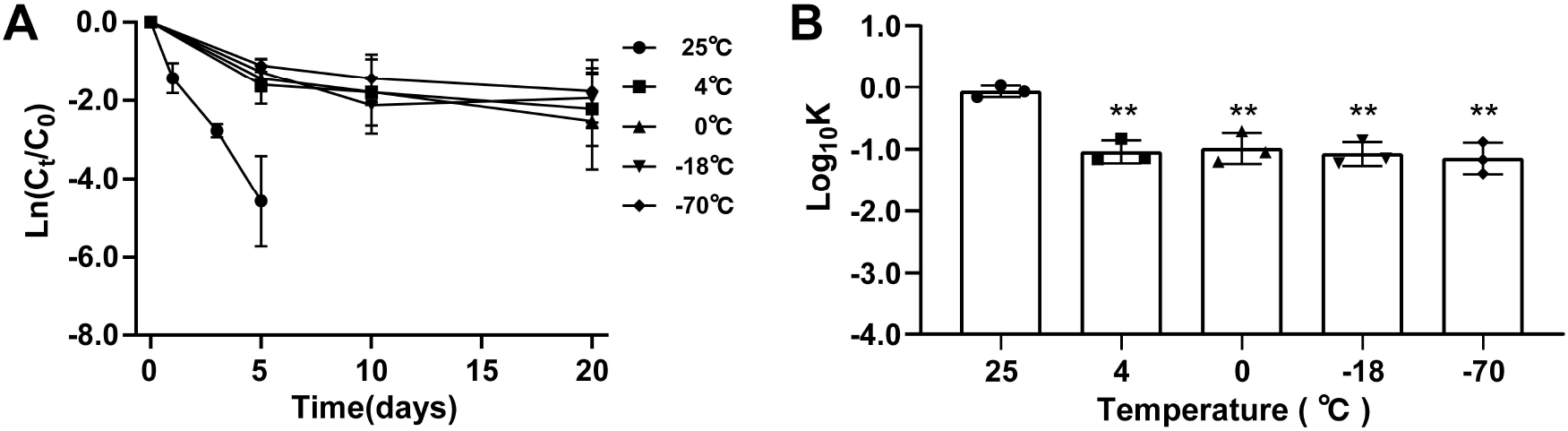
The stability of SARS-CoV-2 pseudovirus under cold-chain temperatures. (A) The decay curves of SARS-CoV-2 pseudovirus on PE plastic over 20 days under different temperatures. (B) The decay rate constants *k* of SARS-CoV-2 pseudovirus on PE plastic under different temperatures. **, *p* <0.01 compared with 25° C.

The pandemics of COVID-19 in the seafood market of different cities suggested that the seawater environment might have the potential to protect SARS-CoV-2 viability. And the SARS-CoV-2 RNA has been frequently detected on the outer packages of the frozen seafood, we further investigated the potential protective effect of seawater on SARS-CoV-2. SARS-CoV-2 pseudovirus was suspended in seawater and deionized water at -18° C and further maintained 8 times freeze-thawing cycles treatment to mimic the loading-unloading process of cold-chain. The stability of SARS-CoV-2 pseudovirus suspended in seawater and deionized water is shown in Fig.3. The decline concentration of SARS-CoV-2 pseudovirus in deionized water was higher than that in seawater under -18° C and repeat freeze-thawing cycles temperature conditions (Fig. 3A and 3B). However, there was no significant difference in the decay rate constants *k* of SARS-CoV-2 pseudovirus between deionized water and seawater under -18° C. However, the decay rate of SARS-CoV-2 pseudovirus in seawater was significantly slower than that in deionized water under the repeat freeze-thawing cycles (Fig. 3C). For the infectivity of SARS-CoV-2 on the suspension medium, the mean decay rate constant *k* ranged from 0.13/day in the seawater to 0.25/day in the deionized water at repeat freeze-thawing cycles with *r*^2^ values from 0.97 to 0.92 (Table 3). Accordingly, seawater but not deionized water protected SARS-CoV-2 pseudovirus thermal stability. we demonstrated that cold-chain temperature protects SARS-CoV-2 pseudovirus survival for a long time and decreases the decay rate of the virus compared to exposure *in vitro* under room temperature. Seawater did not decrease the decay of SARS-CoV-2 pseudovirus *in vitro* at -18° C, but protected the survival and enhanced the thermostability of SARS-CoV-2 pseudovirus *in vitro* under the condition of the repeated freeze-thawing cycle, Researchers have shown that salt enhances the thermostability of viruses by increasing van der Waals force and decreasing repulsive electrostatic force (CRAIG WALLIS; Meister et al., 2020; MELNICK)., and could enhance the risk of the SARS-CoV-2 transmission.

**Table 3.** The decay rate of SARS-CoV-2 pseudovirus under seawater and deionized water at-18° C

| Temperature | *K* (mean ± SD) | [95% CI slope] | R<sup>2</sup> |
| --- | --- | --- | --- |
| Seawater at -18°C | 0.141±0.009 | -0.1595 to -0.1224 | 0.95 |
| Deionized water at -18°C | 0.184±0.020 | -0.2272 to -0.1402 | 0.87 |
| Seawater at freeze-thawing cycles | 0.133±0.011 | -0.1573 to -0.1093 | 0.97 |
| Deionized water at freeze-thawing cycles | 0.2538±0.013 | -0.2813 to -0.2263 | 0.92 |

NS: Non-significant deviation from the model; SD: standard deviation.

**Figure 3.**
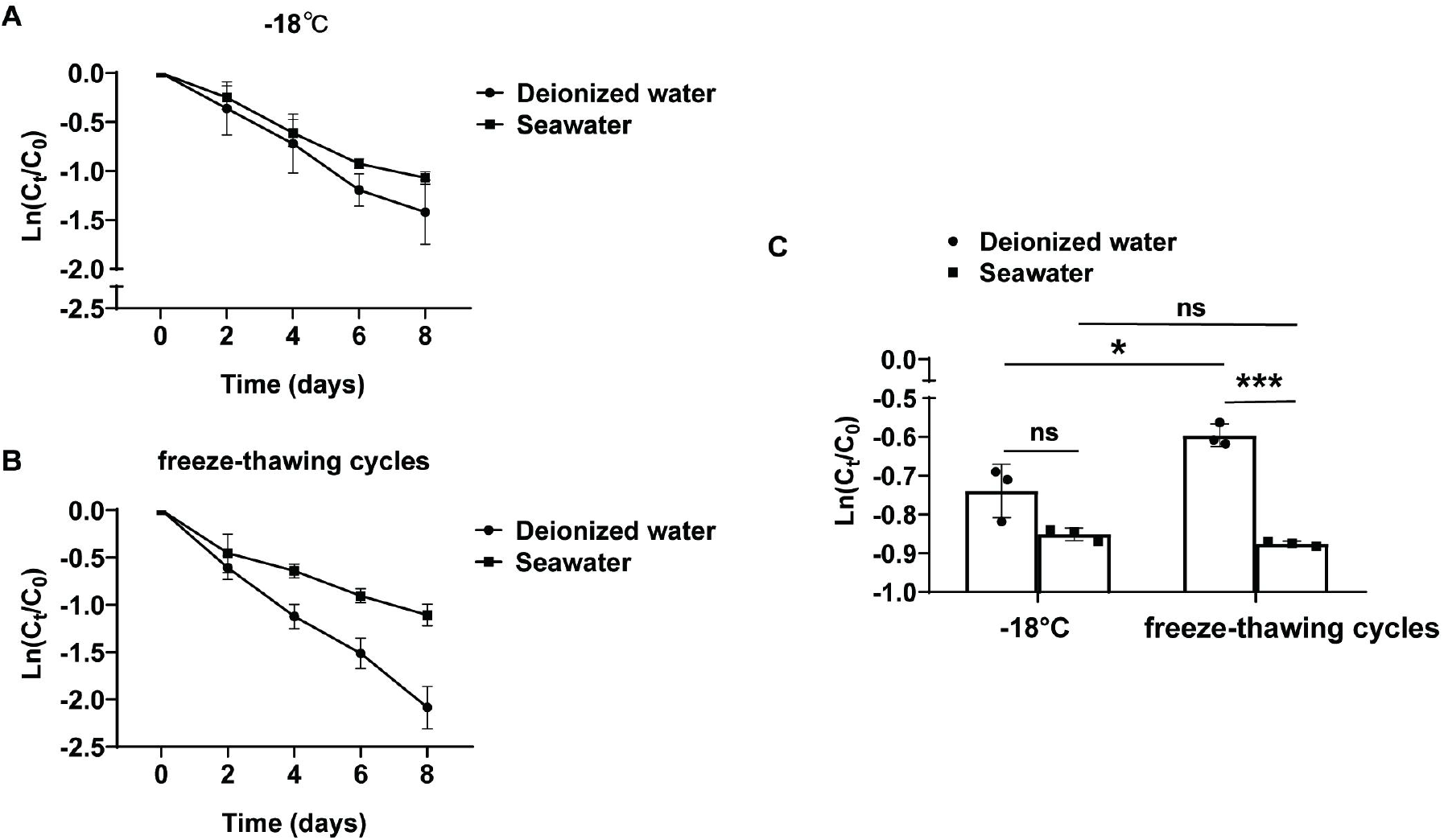
The protective effect of seawater on SARS-CoV-2 pseudovirus at -18° C. The decay curves over time (days) of SARS-CoV-2 pseudovirus, which was solutes in seawater or deionized water, on PE plastic at -18° C (A) and under 8 times freeze-thawing cycles (B). (C) Decay rate constants *k* of SARS-CoV-2 pseudovirus at -18° C and under the condition of freeze-thawing cycles. ns., no significantly difference; *, *p*<0.05; ***, *p*<0.001.

### 3.3 The disinfection potential of LED visible light illumination on SARS-CoV-2 pseudovirus under cold-chain conditions

To explore the possible methods to inactivate SARS-CoV-2 under the cold-chain environment, the LED visible light containing spectral wavelengths from 450 nm to 780 nm was exposed to SARS-CoV-2 pseudovirus contaminated surfaces that are usually used in seafood packing materials, including PE plastic and corrugated carton, which is smoother than cardboard, under -18° C for the indicated time. The concentration of SARS-CoV-2 virus on the PE plastic and corrugated carton declined while exposed to LED visible light compared with the no-light-treatment controls (Fig.4A and 4B). In addition, a total of 2-fold reduction of virus load was observed at 180 min on PE plastic and corrugated cartons using LED visible light compared with the controls. Furthermore, the results of decay rate constants *k* showed that irradiation with LED visible light exposure significantly increased the decay rate of SARS-CoV-2 pseudovirus on PE plastic and corrugated carton under -18° C compared with the no light exposure controls (Fig. 4C and Fig. 4D). For the infectivity of SARS-CoV-2 pseudovirus on LED visible light treatment, mean decay rate constants *k* ranged from 0.004/min at PE plastic to 0.007/min at corrugated carton surfaces. However, in the dark condition, the mean decay rate constants *k* ranged from 0.002/min at PE plastic to 0.003/min at corrugated carton surface (Table 4). To further confirm the survival rate of pseudovirus, we detected the total RNAs of pseudovirus using agarose gel electrophoresis. The results showed that the electrophoresis band of LED visible light-treated pseudovirus RNAs was lighter than those of no-light-treatment controls (Fig. 4E). The quantitative analysis of RNA bands further showed that LED visible light treatment significantly decreased the total RNA of pseudovirus compared with that of no-light-treatment control (Fig.4F). These results suggest that LED visible light can be used as a simple and effective cryogenic physical disinfection tool, which can be applied to reduce the risk of carrying the virus in a cold-chain environment.

**Figure 4.**
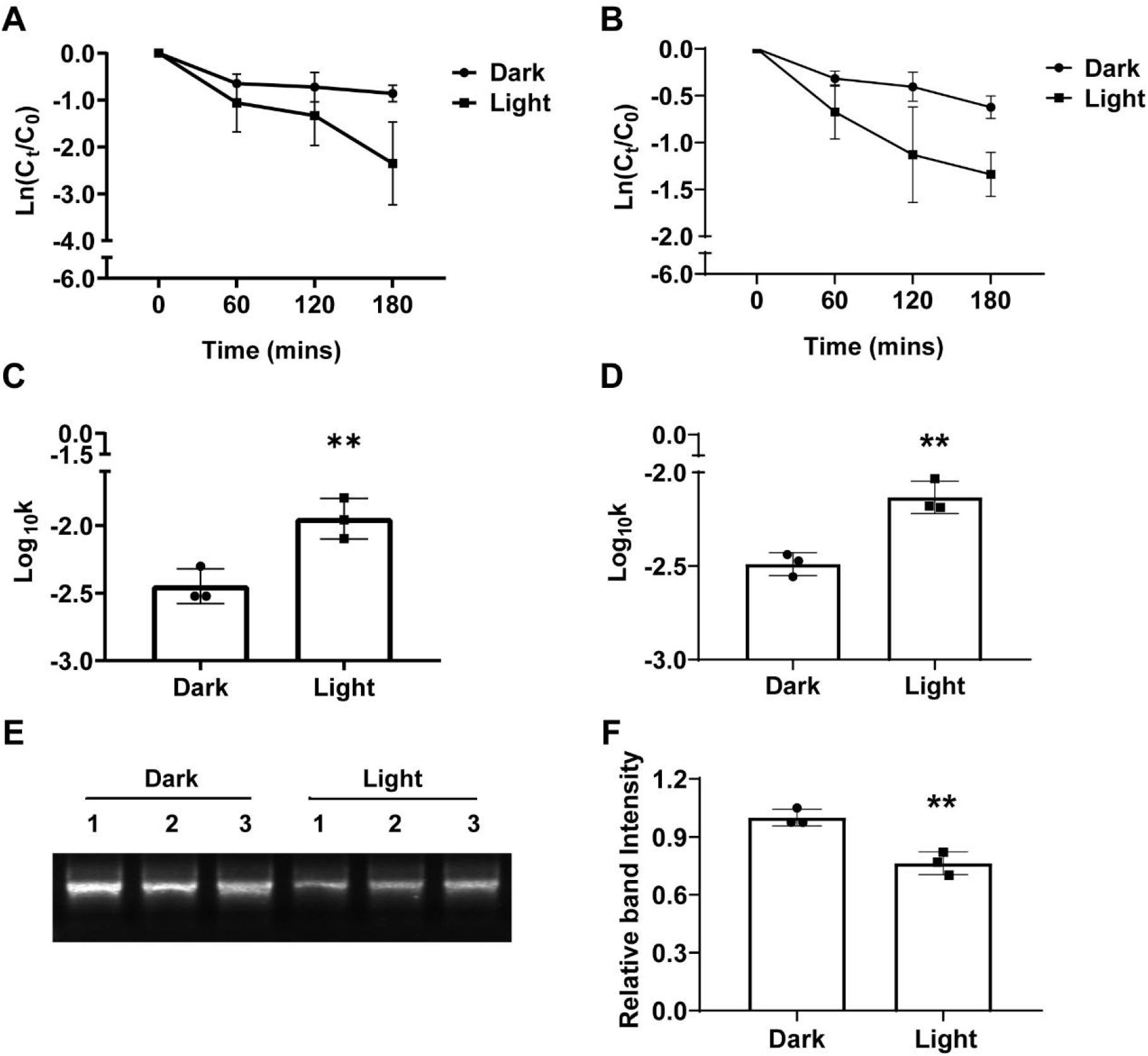
The disinfection potential of LED visible light from 450 nm to 780 nm on SARS-CoV-2 pseudovirus at -18° C. The decay curves over time (min) of SARS-CoV-2 pseudovirus on PE plastic (A) and corrugated carton (B) at -18° C that expose or un-expose to LED visible light (450 nm ∼ 780 nm). The decay rate constants *k* of LED visible light exposed SARS-CoV-2 pseudovirus on the PE plastic (C) and corrugated carton (D) at -18° C were significantly higher than that of light non-exposed control. (E) Genomic RNAs of SARS-CoV-2 pseudovirus on the PE plastic that was exposed or unexposed to LED visible light at -18° C were determined by agarose gel electrophoresis. (F) The RNA band intensities were quantified analysis using gray-scale scanning software. *, *p*<0.05, **, *p*<0.01.

**Table 4.** The decay rate of SARS-CoV-2 pseudovirus under the visible light illumination

| Groups | *K* (mean ± SD) | [95% CI slope] | R<sup>2</sup> |
| --- | --- | --- | --- |
| PE plastic-Darkness | 0.002±0.003 | -0.003 to -0.002 | 0.8684 |
| PE plastic-Light | 0.004±0.005 | -0.005 to -0.002 | 0.6713 |
| Corrugated Carton-Darkness | 0.003±0.001 | -0.004 to -0.002 | 0.8305 |
| Corrugated Carton-Light | 0.007±0.001 | -0.010 to -0.004 | 0.7587 |

NS: Non-significant deviation from the model; SD: standard deviation

### 3.4 The effect of airflow movement on the viability of SARS-CoV-2 pseudovirus under the cold-chain environment

We further analyzed the effect of airflow movement on the viability of SARS-CoV-2 pseudovirus at -18° C. The decay curve over time of SARS-CoV-2 pseudovirus on the PE plastic and corrugated carton under -18° C was decreased while pseudoviruses were exposed under the airflow for 32 h compared with that under a non-airflow movement environment (Fig.5A and 5B). In addition, the decay rate constants *k* of SARS-CoV-2 pseudovirus on the PE plastic and corrugated carton at - 18° C were markedly increased by 1.5-fold in the airflow movement group compared with that in the non-airflow movement group (Fig.5C and 5D). For the infectivity of SARS-CoV-2 on airflow movement treatment, mean decay rate constants *k* ranged from 0.051/h at PE plastic to 0.075/h at corrugated carton surface. However, under non-airflow movement conditions, mean decay rate constants *k* ranged from 0.045/h at PE plastic and 0.064/h at corrugated carton surface (Table 5). These results implied that airflow movement is a facility for the decay of SARS-CoV-2 in the cold-chain environment.

**Figure 5.**
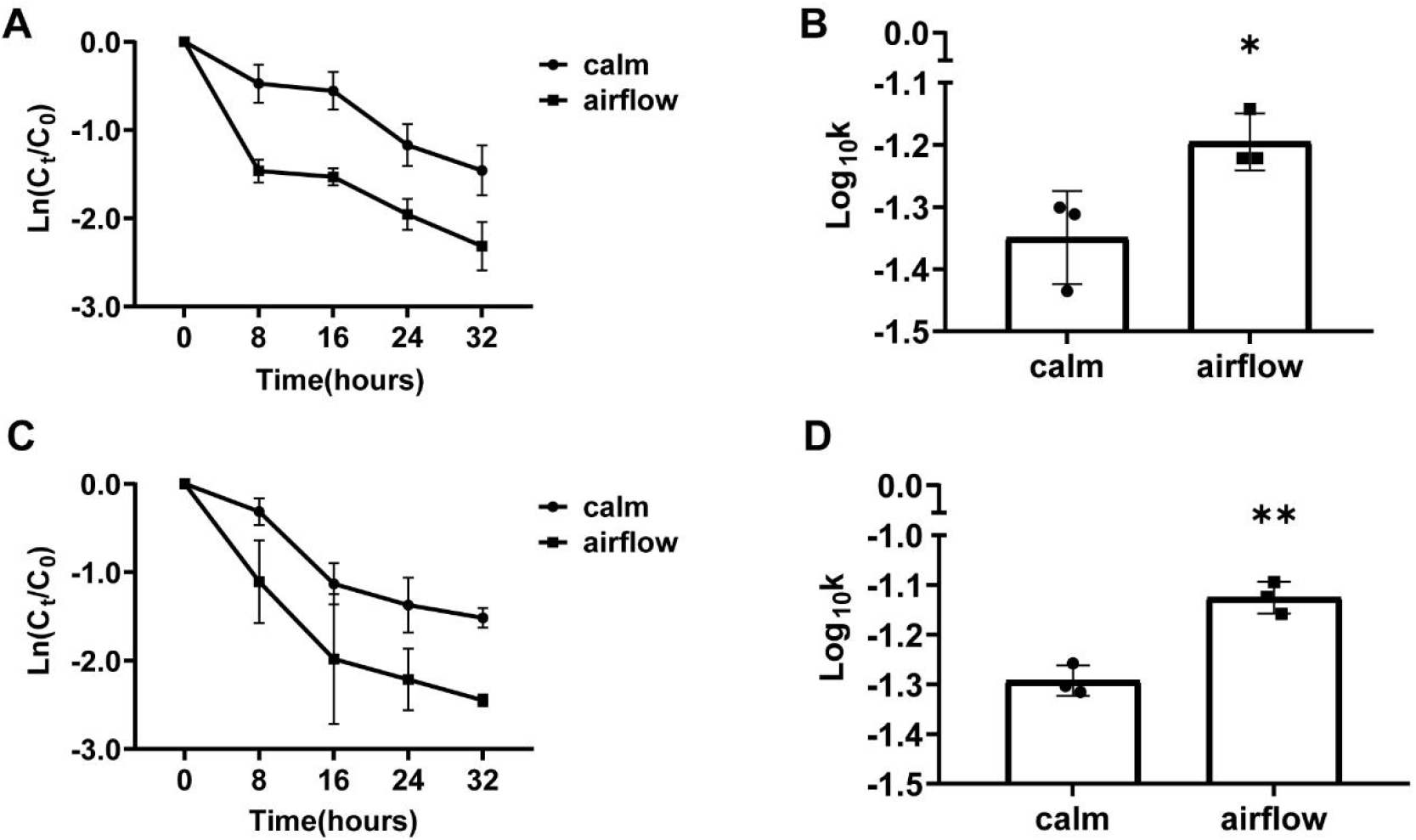
The effect of airflow movement on the viability of SARS-CoV-2 pseudovirus at -18° C. The decay curves over 32 hours of SARS-CoV-2 pseudovirus on PE plastic (A) and corrugated carton (B) at -18° C that was treated or untreated with the airflow movement (3 m/s). The decay rate constants k of wind-treated SARS-CoV-2 pseudovirus on the PE plastic (C) and corrugated carton (D) at -18° C were significantly higher than that of non-airflow movement controls. *, *p*<0.05; **, *p*<0.01.

**Table 5.** The decay rate of SARS-CoV-2 pseudovirus under the air circulation

| Groups | *K* (mean ± SD) | [95% CI slope] | R<sup>2</sup> |
| --- | --- | --- | --- |
| PE plastic-Calm | 0.045±0.005 | -0.056 to -0.035 | 0.87 |
| PE plastic-Wind | 0.064±0.008 | -0.082 to -0.046 | 0.82 |
| Corrugated Carton-Calm | 0.051±0.006 | -0.063 to -0.039 | 0.87 |
| Corrugated Carton-Wind | 0.075±0.011 | -0.099 to -0.051 | 0.80 |

NS: Non-significant deviation from the model; SD: standard deviation

## 4. Discussion

TEM images of negative-stained SARS-CoV-2 particles revealed their pleomorphic spheres with 60-140 nm diameter. The outer membrane of virus particles protruded with quite distinctive spikes that rendered virions a solar corona appearance (Hopfer et al., 2021; Mittal et al., 2020; Zhu et al., 2020). In this study, the particle size and structure SARS-CoV-2 pseudovirus was similar to wild-type virus particles observed by Zhu et al. The infectivity of SARS-CoV-2 is related to the expression level of ACE2 (the receptor of SARS CoV-2) in the target cells. In addition, SARS-CoV-2 virions exhibit longer viability on stainless steel and PE plastic than on copper or cardboard (van Doremalen et al., 2020). Moreover, with the persistence of SARS-CoV-2 on the different surfaces, such as plastic, and stainless steel, this virus is viable on these surfaces for several days (Kampf et al., 2020; Kwon et al., 2021). SARS-CoV-2 decays more rapidly on the porous surfaces than on the non-porous surfaces, but there is no obvious difference in the decay rate of SARS-CoV-2 among non-porous surface types such as PE plastic, stainless steel, or nitrile glove (Biryukov et al., 2020; Chin et al., 2020). Therefore, our findings provide an ideal non-pathogenicity model of SARS-CoV-2 for investigating the infectivity and stability of the SARS-CoV-2 under the cold-chain environment *in vitro*.

Lower environmental temperature correlates with an increased risk of SARS-CoV-2 transmission. Researchers have reported temperature as an important factor affecting the stability and infectivity of the SARS-CoV-2 (Chin et al., 2020). SARS-CoV-2 is highly stable at 4° C with no obvious effect on viability for 14 days (Biryukov et al., 2021; Sharma et al., 2021). Reports from the literature suggested that the cold-chain could maintain the SARS-CoV-2 infectivity and stabilities for days to weeks (Biryukov et al., 2021; Li et al., 2021). In addition, In this study, we demonstrated that cold-chain temperature protects SARS-CoV-2 pseudovirus survival for a long time and decreases the decay rate of the virus compared to exposure *in vitro* under room temperature. Seawater did not decrease the decay of SARS-CoV-2 pseudovirus *in vitro* at -18° C, but protected the survival and enhanced the thermostability of SARS-CoV-2 pseudovirus *in vitro* under the condition of the repeated freeze-thawing cycle.

Researchers indicated that ultraviolet (UV) light, sunlight, 405 nm visible light, and 400 nm to 420 nm standard LED light effectively inactivate SARS-CoV-2 at 20°C (Heilingloh et al., 2020; Rathnasinghe et al., 2021; Ratnesar-Shumate et al., 2020). In this study, we further confirmed that a longer wavelength of LED visible light in the 450 nm to 780 nm range effectively decontaminates and degrades the genomic RNA of SARS-CoV-2 pseudovirus on the PE package surface under a frozen environment. Sunlight irradiation is inconvenient to apply in the closed cold-chain transportation environment and a long time UV light treatment affects the flavor and texture of dried fish fillets and fresh meat (HooD, 1980; Park et al., 2014). Our findings support that illumination with LED visible light may be an ideal decontamination method for the outside of packages in the process of closed cold-chain transportation.

In addition, reports from literature indicated that increased air exchange rate and airflow movement by window-open, air condition or fan significantly reduce the infection risk of COVID-19 in-car and on trains under room temperature (Kumar et al., 2021a; Shinohara et al., 2021). The airflow movement environment accelerates the evaporation of the droplet, changes envelope protein, and then affects virus stability under room temperature (Dbouk et al., 2020; Jayaweera et al., 2020). However, no previous study has evaluated the effect of airflow movement on SARS-CoV-2 viability in a low-temperature environment. This study has demonstrated for the first time that airflow movement reduces SARS-CoV-2 pseudovirus viability under the cold-chain environment.

## 5. Conclusions

In this study, a non-pathogenic SARS-CoV-2 pseudovirus model was provided to confirm the reason for the object-to-human transmission route of COVID-19 and to explore the effective disinfection methods under the cold-chain environment. The research showed the cold-chain temperature and salt as the risk factors that could prolong the SARS-CoV-2 viability and increase the risk of SARS-CoV-2 transmission. Laboratory studies suggested that LED visible light exposure and airflow movement could promote the inactivation of SARS-CoV-2 in the cold-chain environment.

## Acknowledgments

We are grateful to the Institute of Molecular and Clinical Pharmacology, Guangzhou Medical University for the TEM assay. We are grateful to Qian-Qian Zhang for sharing pLV-EGFP-N and psPAX2 plasmid, and Weisheng Guo for giving HEK293T and 293T-hACE-2 cells.

